# Liquid biopsies for omics-based analysis in sentinel mussels

**DOI:** 10.1101/720219

**Authors:** France Caza, Philippine Granger Joly de Boissel, Richard Villemur, Stéphane Betoulle, Yves St-Pierre

## Abstract

Liquid biopsy of plasma is a simple and non-invasive technology that holds great promise in biomedical research. It is based on the analysis of nucleic acid-based biomarkers with predictive potential. In the present work, we have combined this concept with the FTA technology for sentinel mussels. We found that hemocytes collected from liquid biopsies can be readily fixed on FTA cards and used for long-term transriptome analysis. We also showed that liquid biopsy is easily adaptable for metagenomic analysis of bacterial profiles of mussels. We finally provide evidence that liquid biopsies contained circulating cell-free DNA (ccfDNA) which can be used as an easily accessible genomic reservoir. Sampling of FTA-fixed circulating nucleic acids is stable at room temperature and does not necessitate a cold-chain protection. It showed comparable performance to frozen samples and is ideally adapted for sampling in remote areas, most notably in polar regions threatened by anthropogenic activities. From an ethical point of view, this minimally-invasive and non-lethal approach further reduces incidental mortality associated with conventional tissue sampling. This liquid biopsy-based approach should thus facilitate biobanking activities and development of omics-based biomarkers in sentinel mussels to assess the quality of marine ecosystems.

## Introduction

Because of their ability to accumulate xenobiotics in their tissues, their wide distribution, and their ecological and economical importance, blue mussels (*Mytilus* spp.) have long been recognized as good biological indicators for monitoring the effects of pollution and climate change in marine habitats [1]. Not surprisingly, a relatively large number of biomarkers have been developed, often using a multi-tier approach. They include functional biomarkers based on enzymatic activities or cellular functions, or on measuring concentrations of specific stress indicators [2–4]. In most cases, measure of these biomarkers requires on-site analysis or tissue biopsy performed by trained personnel. It also often requires an uninterrupted cold chain to maintain sample quality. For sampling in remote areas, such as polar regions, this implies complex logistical challenges and considerable risk and cost associated with transportation and storage. The use of formalin-fixed tissue biopsy is an alternative, although formalin is known to affect DNA quality. There are, however, novel methods for collection, storage and shipping of biological samples that offers some solutions to these problems. The Flinders Technology Associates (FTA) cards, for example, can be used to fix and preserve nucleic acids for long-term periods at ambient temperature with high stability [5,6]. This approach has been particularly useful for identification/genotyping and long-term surveillance of infectious pathogens in remote areas at reduced shipping costs [7–9]. The use of FTA cards for monitoring marine habitat with mussels as sentinel species, however, will require concomitant development of nucleic acid-based biomarkers. Fortunately, immense progress in the development has been achieved thanks to new methods in gene sequencing and the need for accurate predictive biomarkers in medicine. Nucleic acid-based biomarkers are particularly well-adapted for liquid biopsies. Liquid biopsy is a concept-based approach that is gaining significant attention in the biomedical area. The concept is that a clinician is able to use a small sample of blood collected from a patient to obtain vital genetic information that can be used: 1) to predict the onset of a disease, 2) to assess the severity of a disease (prognostic), 3) to evaluate and measure with high precision the efficacy of a given treatment, and 4) to determine long-term survival and relapse. It is designed to replace the use of logistically complicated, high cost invasive tissue biopsies to assess ongoing tissue damage. It is largely based on the analysis of circulating cell-free DNA (ccfDNA), which is considered as a genomic reservoir that is easily accessible for studying genetic traits, including genomic alterations [10, 11]. Recent studies in humans have further shown that ccfDNA can be used to identify damages in specific tissues [12]. Liquid biopsies are also compatible for metagenomic data analysis. Microbiome information is now considered an important component of the biomarker schematic. With the help of recent advances in next generation sequencing, microbiome-associated biomarkers, just like ccfDNA, can be used in clinical practice for risk prediction, diagnosis and progression of a disease, or to predict and modulate response to treatment [13, 14]. In marine ecology, microbial biomarkers are also being used for measuring the impact of anthropogenic activities on marine ecosystems [15, 16]. In the present work, we have combined the concept of liquid biopsy and the use of FTA cards to develop a liquid biopsy-based sampling method that involves fixation of nucleic acids on FTA cards. This method is simple, does not require maintaining a cold chain and is compatible with transcriptome analyses. We have also identified, for the first time, circulating cell-free ccfDNA in hemolymph of mussels and performed 16SrDNA gene-based bacterial microbiota profiles of mussels using FTA-fixed bacterial DNA collected from liquid biopsies.

## Material and Methods

### Liquid biopsy

Adult specimens (55–70 mm length) of blue mussels *Mytilus spp.* were obtained commercially in Canada and placed in a temperature-controlled (4°C) aerated aquarium containing 10 L of 32‰ artificial saline water (Reef Crystal artificial marine salt, Instant Ocean, VA, USA). For *M. desolationis* and *Aulacomya ater* (*A. ater*) specimens, they were collected on the intertidal rocky shore of Port-aux-Français (Kerguelen Islands, France) as previously described [17]. In all cases, hemolymph was collected from single individuals, without pooling. For routine preparation of liquid biopsies on FTA cards, the intervalvar liquid was removed with the tip of a knife and hemolymph was withdrawn from the adductor muscle using a syringe fitted with a 25 gauge needle. Samples (<1ml of hemolymph) were immediately transferred into sterile 1.5 ml Eppendorf tube and centrifuged for 3 minutes at maximum speed (approx. 3000 x *g*) at room temperature using a battery-powered mini-centrifuge (TOMY, Japan) (**Fig 1**). After centrifugation, aliquots (50 μl) of supernatants were spotted on individual discs (10 mm diameter) on Whatman 903™ FTA cards (Whatman, Z694827, Sigma-Aldrich, Oakville, ON, Canada). In some experiments, Whatman FTA Gene Cards (GE Healthcare, Marlborough, MA) were used. The remaining supernatant was discarded and hemocyte pellets gently resuspended and spotted on FTA cards. The cards were dried for 15 min at room temperature and stored individually in small (3” x 4”) clear 2 mil plastic ziplock baggies containing one silica gel desiccant (1gr/bag) moisture absorber. When indicated, hemolymph were pooled, transferred into a sterile 50 mL and centrifuged for 15 min at 3000 x *g* on a table low speed centrifuge (Beckman Coulter Allegra 6R). Unless otherwise indicated, all FTA cards were kept at room temperature in the dark.

**Fig 1.**
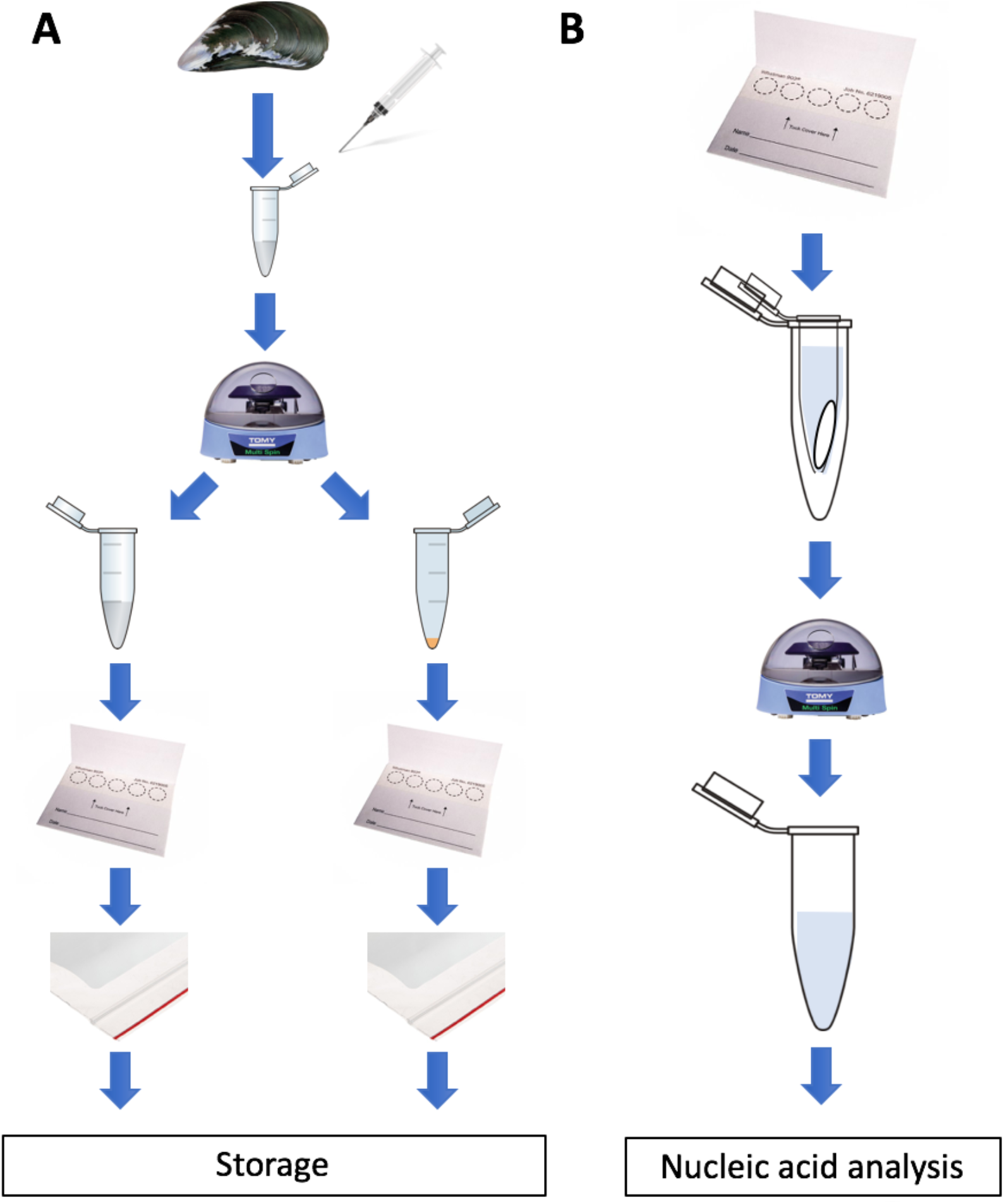
Diagrammatic summary of the sampling protocol. *Left*, for routine preparation of liquid biopsies on FTA cards, hemolymph was withdrawn from the adductor muscle. Samples (approx. 1ml of hemolymph) were immediately transferred into sterile 1.5 ml and centrifuged at room temperature using a battery-powered mini-centrifuge (TOMY, Japan). After centrifugation, aliquots (50 μl) of supernatants were spotted on individual discs (10 mm diameter) on FTA® cards. The remaining supernatant was discarded and hemocyte pellets gently resuspended and spotted on FTA cards. The cards were dried for 15 min at room temperature and stored individually in small plastic Ziploc baggies containing a silica gel desiccant moisture absorber. *Right*, to recover nucleic acids, discs were carefully placed at the bottom of a sterile microcentrifuge tube with a hole punctured in its bottom. The tube was then transferred into a larger sterile microcentrifuge tube before isolation of nucleic acids as described in details in the material and methods section.

### Isolation of ccfDNA from hemolymph

ccfDNA from hemolymph (5 ml) was carried out using the NucleoSpin Plasma XS kit (Macherey Nagel) with optimized manufacturer’s protocols. Purified DNA was quantified by the PicoGreen assay according to the manufacturer’s recommendations [18].

### Isolation of nucleic acid

For RNA isolation and purification, individual 12 mm^2^ disks from FTA cards were carefully placed at the bottom of a sterile 0.6 ml microcentrifuge tube with a hole punctured in its bottom by a 18G needle. The tube was then transferred into a sterile 1.5 ml microcentrifuge tube before addition of TRIzol (800 μl) to the upper tube. The cards were soaked with TRIzol for 10 min at room temperature and centrifuged for 3 min at high speed (8000 rpm) in a table centrifuge (BioFuge Fresco, Heraeus Instrument, Grimsby, ON, Canada). The TRIzol solution was transferred and topped with 0.2 ml of chloroform. Tubes were incubated 3 min at room temperature and centrifuged at 12000 x *g* for 15 min at 4°C. The aqueous phase was used to perform RNA extraction following a standard protocol. The RNA was dissolved in RNase/DNase free water and quantified using the Thermo Scientific™ NanoDrop™ spectrophotometer. DNA was isolated using an adapted the prepGEM™ Universal kit for Blood storage cards (Zygem, Charlottesville, VA). Briefly, 3 mm disc were placed on an eppendorf tube in washed with 100 μl of DNA-free water and DNA recovered with the prepGEM solution following 5 min incubations at 75°C and 95°C. Nucleic acid samples were stored at −20°C until further analysis. Purified DNA was quantified by the PicoGreen assay (Lumiprobe, Hunt Valley, MD) according to the manufacturer’s recommendations [18].

### RT-PCR analysis

Total cellular RNA was isolated from cell pellets fixed on FTA cards using the TRIzol reagent (Life Technologies) as described above. First-strand cDNA was prepared from 2 μg of cellular RNA in a total reaction volume of 20 μl using the High Capacity cDNA Reverse Transcription Kit from Thermo-Fisher. qPCR specificity was carried out with a Rotor-Gene® 3000 (Corbett Research) with primers listed in **Table I**. All qPCR reactions were carried out in 20 μl with the QuantiFast SYBR Green PCR Kit (Qiagen). qPCRs were amplified using the following conditions: 95°C for 5 min, followed by 40 cycles at 95°C for 10 s and 60°C for 30 s with a single acquisition. A final melting curve analysis was performed using at 95°C for 5 s, 65°C for 60 s and 97°C with a continuous acquisition. To ensure primer specificity, melting temperature curves were determined for each amplicon. PCR products were run on a 1.5% agarose gel to visualize the bands and ensure a unique band is amplified. The expression level (log_2_ of the fold change) was expressed 1) calculating the delta CT (∆CT) by subtracting the averaged threshold cycle (CT) value of the three housekeeping genes (*EF1γ, EF1α* and *SNX14*) from the CT value of the target gene and 2) comparing the fold change of non-exposed mussels versus thermal-exposed mussels [19]. In some experiments, first-strand cDNA was prepared directly from 3 mm discs obtained using a Harris Uni-core punch. In this case, the discs were washed for 5 min at room T°C with 200 μl of Whatman FTA solution. The discs were then dried before adding RT reaction reagents. A volume of 5 μl was then trasnferred into another eppendorf tube for PCR amplification, as described above.

**Table I.**
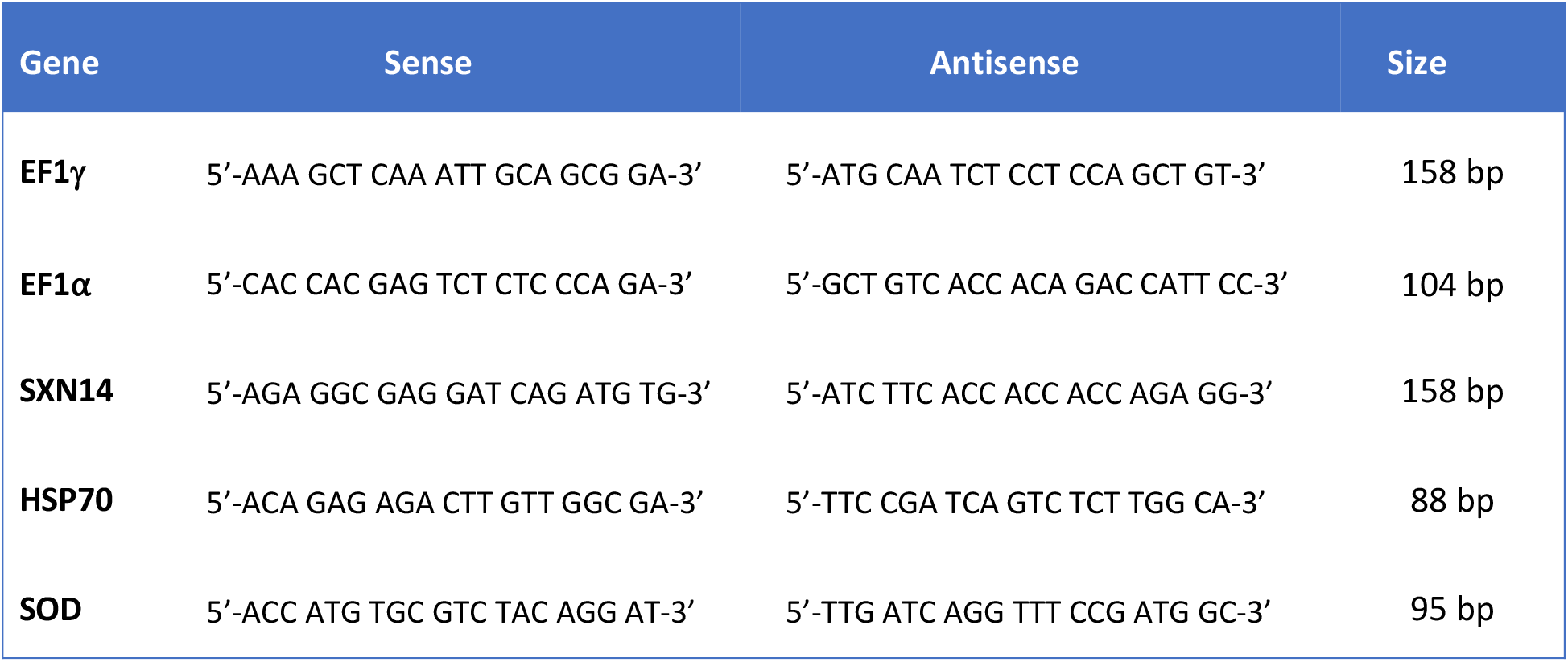
List of primers.

### Thermal stress

Following acclimatization, groups of ten mussels of similar length were transferred into tank containing oxygenated seawater at 30°C or at 4°C (control). After 30 min, liquid biopsies were collected. Hemocytes pellets and hemolymph were either fixed on FTA cards or kept frozen at −20°C.

### Taxonomic analysis of hemolymph microbiota

Following fixation on FTA cards, DNA was extracted from hemocyte pellets, as described above. The whole 16S bacterial DNA V2-V3 region was targeted by the primers 28F-519R primers and pyrosequenced by the 454 FLX Roche technologies at Research&Testing Laboratory (http://www.researchandtesting.com/, Texas, USA). An average of 3,000 sequences was generated per sample. A detailed description of bioinformatic filters that were used for analysis can be found at http://www.rtlgenomics.com/docs/Data_Analysis_Methodology.pdf. The metagenomic data from this publication have been deposited to the ENA database http://www.ebi.ac.uk/ena and assigned the identifier PRJEB19465.

### Statistics

All datasets were validated for normality (with Shapiro-Wilk test) and homoscedasticity (with Bartlett and F of Fisher test). Student’s t-test was used as parametric test to compare data set with two groups and a two-way analysis of variance (ANOVA) was used to compare datasets with more than two groups. For non-parametric data, Kruskall Wallis test was used to compare datasets. All data are expressed as the mean *±* S.D. Results were considered statistically significant at P≤0.05.

## Results

### Isolation and analysis of RNA from hemocyte pellets fixed on FTA cards

Because hemocytes are implicated in several physiological functions, such as phagocytosis of pathogens, homeostasis, and reproduction, there is increasing interest in studying how their transcriptome is modulated by environmental stress factors [20–24]. We thus developed a general framewok procedure for collecting hemolymph samples using FTA cards without the need of a cold chain is shown and detailed in **Fig 1a**. Other than a battery-operated centrifugre, it does not require specialized equipment and can be easily incorporated into a small, light-weight portable unit (**S1 Fig**).

To test whether such sampling framework is compatible for multi-omics analysis of hemolymph samples, we first compared RNA yields obtained from frozen hemocytes and hemocytes that were fixed on FTA cards. Isolation of RNA from FTA discs was carried out using a simple TRIzol-based protocol which consisted of soaking FTA discs in TRIzol for 10 min at room temperature (**Fig 1b**). Nucleic acids were then eluted from the discs by brief centrifugation and RNA purified using standard phase separation with chloroform and subsequent precipitation with isopropyl alcohol [25]. Our results showed that similar amount of RNA was recovered from pellets fixed on FTA discs as compared to pellets frozen at −20°C (**Fig 2a**). Both methods yielded similar ratio of the absorbance at 260 and 280 nm. RNA recovery from FTA discs could also be achieved using a commonly-used silica-based RNA extraction kit, which resulted in lower yield compared to the TRIzol-based method but provided purer RNA (**Fig 2b**). We also found that the RNA yield from hemocyte pellets fixed on FTA was not significantly affected by long term storage (up to 6 months) at 4°C or ambient temperatures (**Fig 2c**). RT-qPCR analysis using frozen RNA and RNA fixed on FTA cards showed that both conservation methods performed equally when measuring expression of three commonly used housekeeping genes (*EF1α, EF1γ*, and *SNX14*) (**Fig 3a**). Both methods were also equally efficient measuring the *de novo* expression of stress-related genes (*Hsp70* and *SOD*) in hemocytes collected from mussels subjected to an acute thermal stress (**Fig 3b**).

**Fig 2.**
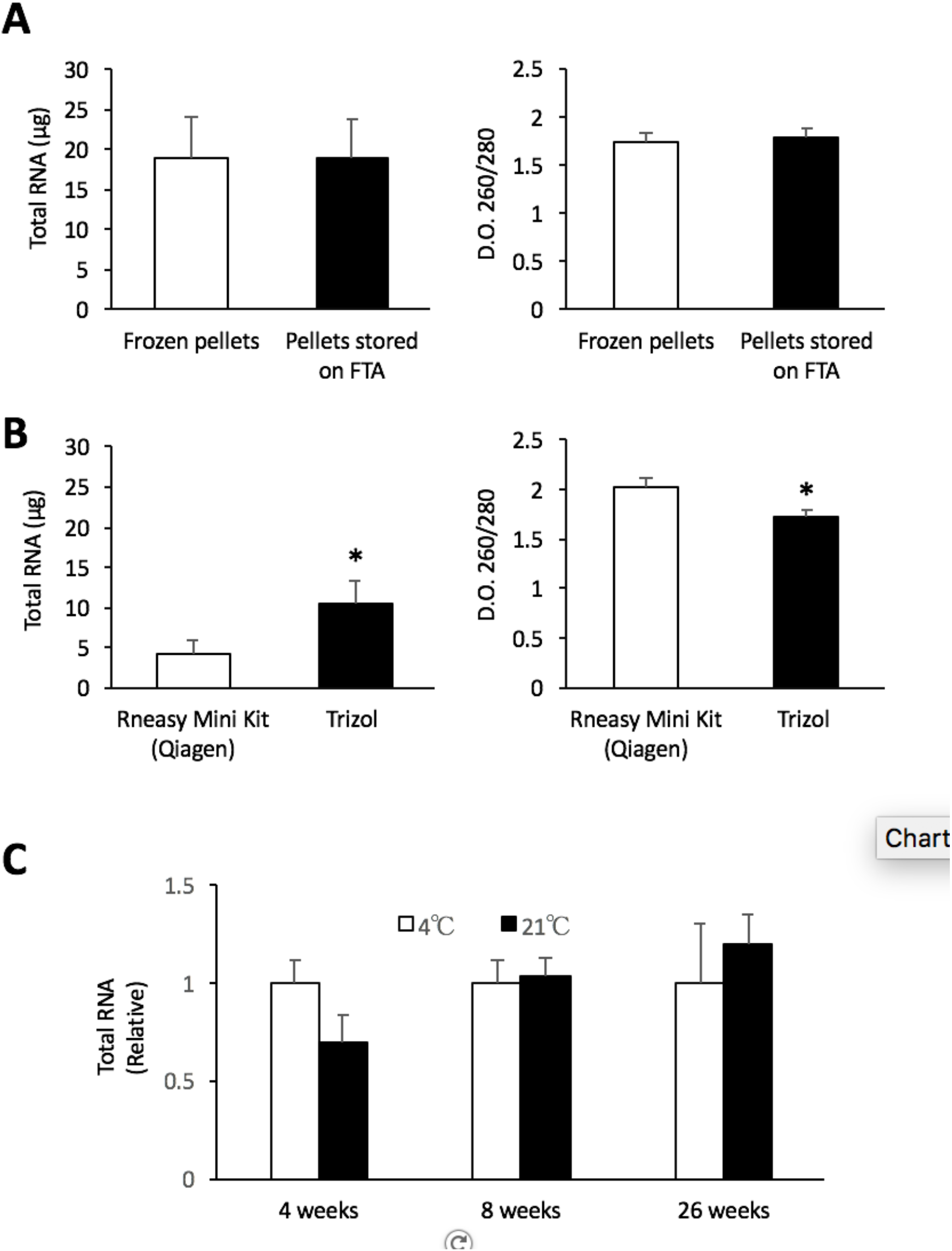
RNA extraction from FTA cards. (**A**) Recovery of total RNA from hemocytes stored frozen at −20°C or on FTA cards using a TRIzol-based protocol. (**B**) Comparative RNA recovery using TRIzol or the RNeasy kit. (**C**) Stability of RNA fixed on FTA cards and kept at different temperatures. * *p* < 0.05. Values are expressed as the total amount of RNA recovered from a 10 mm diameter sample disc FTA cards. No significant (N.S.) differences were found using both methods. Data represent the mean (± *standard deviation*) of two independent experiments, each performed in triplicates.

**Fig 3.**
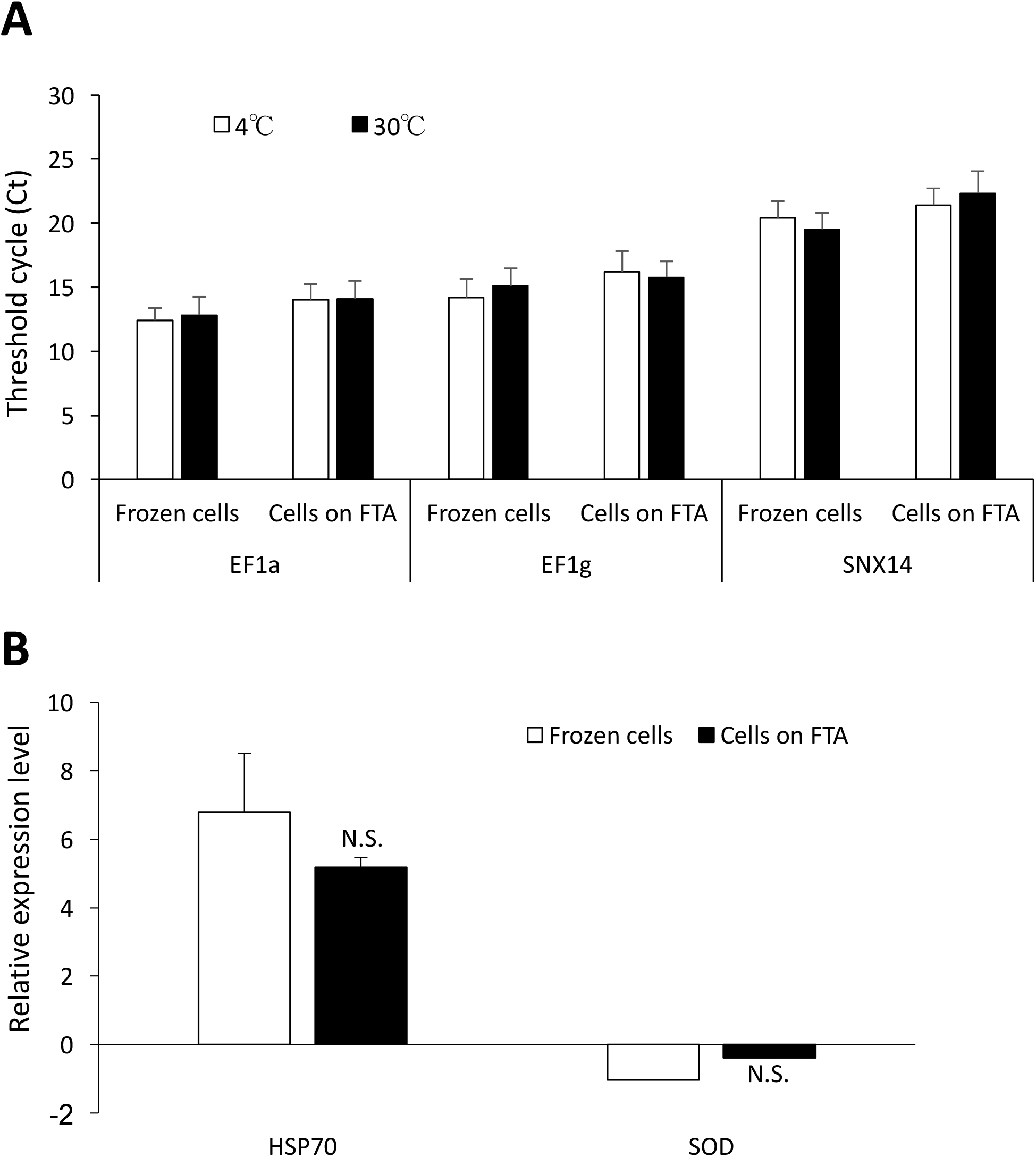
RT-qPCR analysis using frozen and FTA-fixed RNA. (**A**) Cycle tresholds (Ct) values obtained for expression analysis of three housekeeping genes in hemocytes pellets that were either kept frozen or fixed on FTA cards. (**B**) Relative gene expression of *Hsp70* and superoxide dismutase (*SOD*) in hemocytes following an acute thermal stress at 30°C for 30 min. Expression levels are relative to controls (4°C). No significant (N.S.) differences were found using both methods. Data represent the mean (± *standard deviation*) of two independent experiments, each performed in triplicates.

### Bacterial microbiome analysis

Bacteria microbiome-derived profiles are emerging as potent biomarkers not only in the biomedical field, but also in monitoring environmental stress in a vast number of species, including marine organisms [26, 27]. For these experiments, we compared the 16SrDNA gene-based bacterial profiles from frozen and FTA-conserved DNA. Our results showed that both methods generated similar bacterial profiles with regards to phylum (**Fig 4a**). We found no significant differences in genus-level bacterial groups that were most frequently found in hemolymph (**Fig 4b**). The most common bacteria found in the hemolymph was from the genus *Colwellia*. This genius is commonly found in marine habitats, including the hemolymph of blue mussels [16]. Taken together, these findings show that FTA-fixed DNA is perfectably compatible with metagenomic analysis of circulating microbiota.

**Fig 4.**
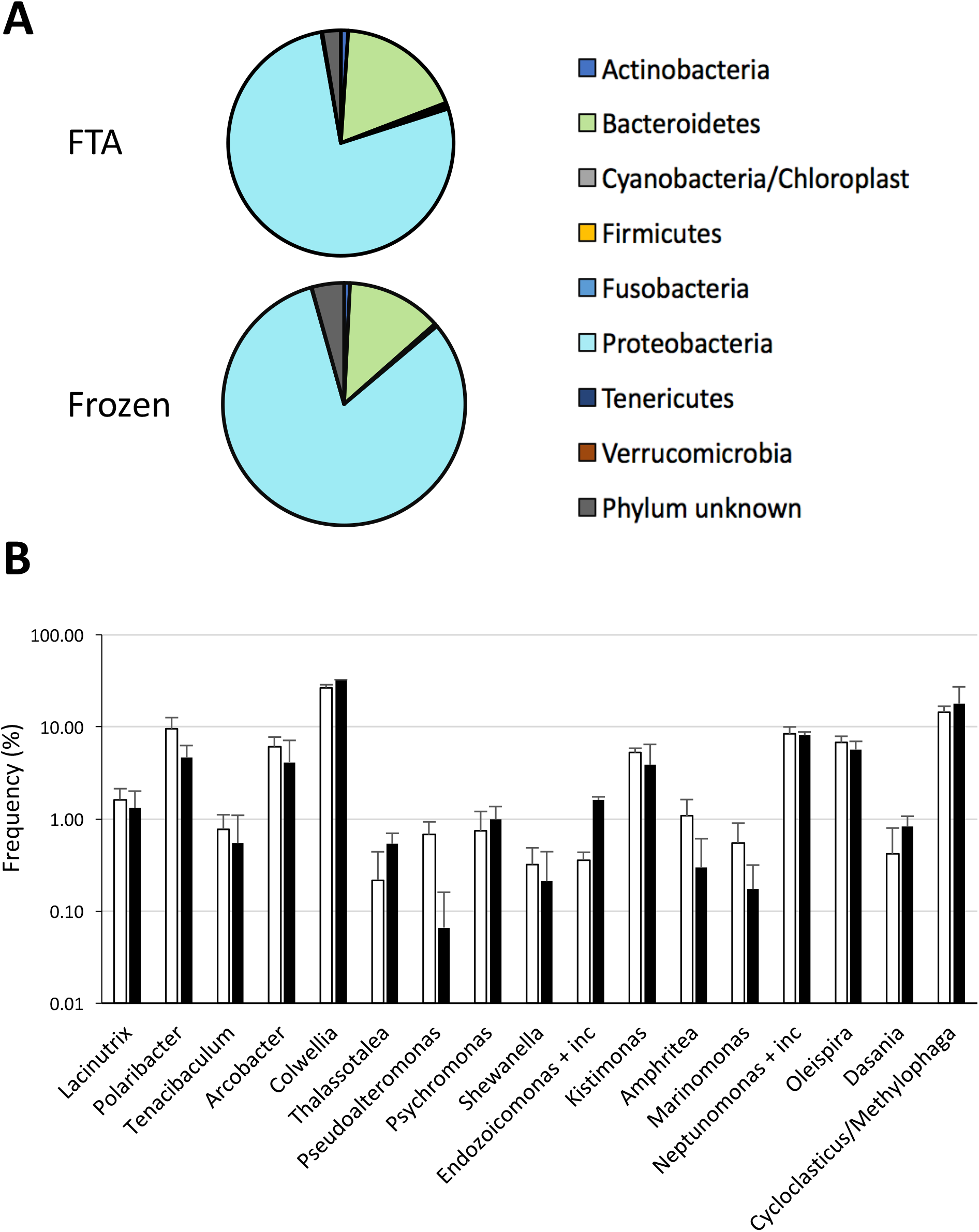
Bacterial profiles of hemolymph. **(A)** Pie chart representation of the relative abundance 16SrRNA of the most common phyla found in liquid biopsies of *Mytilus spp*. **(B)** Bacterial genus differentially abundant in frozen and FTA-fixed liquid biopsies.

### Circulating cell-free DNA

Analysis of ccfDNA from liquid biopsies is another promising avenue for the development of predictive biomarkers. In humans, it serves as a genomic reservoir that is easily accessible with a minimally-invasive procedure. To our knowledge, however, the existence of ccfDNA in hemolymph of invertebrates, and in mussels in particular, has not been reported. Using the same method that is commonly used for isolating ccfDNA (see *material and methods* section for details) from human plasma, we have thus investigated the presence of ccfDNA in mussels. Our results revealed that we indeed found ccfDNA in hemolymph of *Mytilus spp.* (**Table II**) at in a range of concentrations (nanograms/ml) that were similar to that found in humans [28]. Similar results were obtained for *M. desolationis* and *Aulacomya ater* (*A. ater*). To validate the integrity of the ccfDNA fragments in liquid biopsies (cell-free hemolymph) fixed on FTA cards, we carried out real-time qPCR analysis using primers that generated small amplicons and that were specific for *EF1γ*, HSP70 and *SNX14* genes [29] (**Table II**). Our results showed that we could indeed find evidence of ccfDNA fragments encoding *SNX14* and *HSP70 genes* but not *EF1γ*, indicating that ccfDNA fragments are generated is gene-specific, as observed in humans (**Table III**). These results reveal for the first time the presence of ccfDNA in mussels and open the door for its use as a biomarker in liquid biopsies collected from mussels.

**Table II.** Concentration of ccfDNA.

| Samples | ccfdNA (ng/ml) |
| --- | --- |
| Human plasma | 8,72 ± 1,43 |
| *M. trossulus* | 4,23 ± 0,18 |

**Table III.** qPCR analysis of ccfDNA fixed on FTA cards.

| Gene | Template | CT | MT | % of detection |
| --- | --- | --- | --- | --- |
| EF1γ | ccfdNA | 25,31 ± 0,85 | 78,50 ± 1,71 | 0% |
| | NTC | 25,79 ± 0,67 | 79,65 ± 0,21 | |
| SNX14 | ccfdNA | 24,64 ± 2,44* | 83,84 ± 0,47 | 56% |
| | NTC | 31,79 ± 0.01 | N.D. | |
| HSP70 | ccfdNA | 21,25 ± 1,55 * | 81,51 ± 0,14 * | 89% |
| | NTC | 32,16 ± 0 | 80 ± 0 | |

NTC, No template control. CT, threshold cycle, MT, melting temperature, and % of amplicon detection obtained for EF1γ , SNX14 and HSP70 (n=9 mussels, with 3 replicates for each sample. ND = no detected. \* $p < 0.05$ ).

## Discussion

Given the growing concerns over the anthropogenic impact on marine ecosystems, it is increasingly critical to develop biomarkers that can be used as diagnostic and predictive tools, as well as for monitoring the success of the remediation efforts. The use of multi-omics biomarkers is a step in that direction. It obliges us, however, to go back to the drawing board to re-examine our sampling strategy not only for logistical reasons, but for economical reasons as well. This is particularly true for polar regions where the logistic complexity of tissue sampling is a major obstacle. In the present work, we have combined the use of FTA cards and the concept of liquid biopsies for the development of novel omics-based biomarkers in mussels, a widely used sentinel species. More specifically, we have shown that: 1) hemocytes pellets can be readily fixed on FTA cards and used for gene expression analysis; 2) FTA-fixed cell pellets are stable for long term periods when stored at room temperature, facilitating the biobanking activities and shipment to third parties at lower cost when compared to frozen samples; 3) our sampling strategy is easily adaptable for microbiome analysis of bacterial profiles, and 4) ccfDNA is present in mussel hemolymph and can be used as a genomic reservoir for the development of novel biomarkers. Taken together, this study lays the foundation for a logistically-friendly, omics-based multi-biomarker approach using mussels to assess the quality of marine ecosystems.

Until now, the concept of liquid biopsies has been almost exclusively used for guiding clinical practice, especially in oncology, prenatal screening, transplantation and presence of non-human ccfDNA (virus-derived sequences, for example) [10, 30, 31]. One of the main benefits of non-invasive liquid biopsies is that it overcomes many of the drawbacks associated with tissue biopsies. It also provides an accurate snapshot of the genomic landscape, bypassing many problems associated with tissue heterogeneity. In fact, to our knowledge, this is the first report of the presence of ccfDNA in invertebrates. In humans, the normal plasma concentration of ccfDNA is generally below 10-20 ng/ml, which is relatively similar to what we found in hemolymph of mussels. while it can increase by 5–10 times in patients with malignant disease or stress conditions [30]. Future studies will be needed to determine whether measuring ccfDNA levels in mussels can be used as a rapid stress biomarker as it does in humans [28, 32, 33]. Analysis of ccfDNA is also perfectly adaptable for serial samplings of DNA abnormalities, gene expression analysis, point mutations, chromosome translocations, miRegulome, epigenetic changes, etc. For example, analysis of ccfDNA can theoretically be used for measuring mutations in genes associated with hematopoietic neoplasia (such as *p53*), for rapid identification of mussel species, or for any genomic traits. It could also be used to detect the presence of parasitic and viral DNA to reduce the risk for public human health. From an ethical point of view, it reduces incidental mortality associated with conventional tissue biopsies. To develop the full potential of this approach, we are currently using next-generation sequencing to perform a genomic analysis of the entire ccfDNA found in different mussel populations.

Another benefit of combining both FTA-based technology and liquid biopsy is its compatibility with microbiome analysis. In humans, although blood is normally considered a sterile environment, there is increasing studies showing the circulating cells harbor a rich bacterial microbiome that can be used as disease biomarkers [34, 35]. This is also true for bivalves. The microbiome of hemolymph samples collected from the adductor muscle from Pacific oysters, for example, has been shown to undergo significant changes following abiotic and biotic stress factors [36]. A recent study showing that bacterial diversity and richness in mussels were higher in the hemolymph compared to other tissues support the idea that liquid biopsies is a valid approach for microbiome analysis in mussels [16]. Our results further showed that the bacterial profiles obtained from DNA of cells fixed on FTA was not significantly different from that obtained from frozen cells. We found no significant differences in those bacteria that are commonly found in marine ecosystems, such as *Alteromonadales* or *Oceanospirillales* [37, 38]. In fact, our results showed that DNA from cells fixed on FTA cards was more prone to detect those rare bacterial DNA, consistent with the ability of the chemical matrix of FTA cards to stabilize and protect nucleic acids from environmental degradation. Not surprisingly, one of the most abundant bacterial DNA that we found in both FTA cards and frozen samples derived from the *Polaribacter* genus, which species are almost always found in polar or marine environments [39]. The other most frequently found genus were *Colwellia*, *Oleispira*, *Cycloclastricus* and *Neptunomonas*. These bacteria are well known to degrade polycyclic aromatic hydrocarbons (PAH) or short-chain alkans. *Cycloclasticus* bacterial DNA has been recently found in *Bathymodiolus heckerae* mussels where they have established a symbiosis at a 3000m depth in oil-rich habitats near asphalt volcanoes in the Gulf of Mexico [40]. They have also been shown to bloom in marine habitats following the Deep Horizon oil spill [40]. Interestingly, the *Oleispira* genus, another bacterium that has bloomed following the Deep Horizon oil spill, was also among the most frequently found bacterium in *M. edulis*. The third most frequent bacteria found was from the genus *Neptunomonas*, another bacterium implicated in degradation of PAHs and normally found in contaminated sediments [41]. Whether the presence of this microbiota reflects a symbiotic relationship with *M. edulis* is unclear at the present time. Overall, our findings indicate that microbiome analysis in liquid biopsies collected from mussels is an interesting avenue to study dysbiosis in mussels following exposure to environmental stress.

Our study showed that combining liquid biopsies and the FTA technology could be useful for long-term biobanking and retesting of omics biomarkers collected from sentinel mussels, particularly in remote regions. This approach is also economically viable, especially when using the Whatman 903™ FTA cards. Although we have used for this study 10 mm diameter circular FTA discs, we obtained similar RNA yield using smaller discs (**S2a Fig).**If necessary, one could use the the Whatman FTA Gene Cards, which have been adapted for cature on cucleic acids. Although more expensive than regular Whatman 903™ cards, we found that they are more performant for DNA recovery (**S2b Fig**). It is important to note, however, that there are multiple options for isolating nucleic acid from FTA cards. In our hands, we have found that TRIzol was an adequate, low cost, and versatile method for routine RT-qPCR analysis. It can also be combined with commercial kits to obtain purer RNA preparations (**S1 Table**). TRIzol is also suitable for isolation of small RNAs (<200 nucleotides) as compared to the silica-based methods which do not retain small RNAs [42]. This has to be taken into consideration for future studies aimed at testing the possibility of using non-coding RNAs as putative biomarkers in hemolymph of mussels. In humans, non-coding RNAs have shown great promise as potential cancer biomarkers for liquid biopsies. For some experiments, however, we found that RNA does not need to be extracted from the FTA cards. First-strand cDNA templates can be generated directly from a 3 mm punch and subsequently used for standard PCR amplification (**S2c Fig.**)

Another clear benefit of combining liquid biopsy and the FTA technology is from a logistically-point of view. We can now collect samples at remote site with a minimal amount of equipment. Moreover, this approach addresses increasing concerns in field work ethics [43]. Liquid biopsies require sampling of a very small volume of hemolymph and does not lead to animal death. We believe that such approach could also be extended to other marine organisms and offers many advantages for the development of long-term ecological observatories in polar regions threatened by anthropogenic activities.

## Acknowledgments

The authors would like to thank Ms. Marlène Fortier for her technical help, all the personnel from the French Polar Institute Paul Emile Victor (IPEV) and the Terres Australes et Antarctiques Françaises (TAAF) for their help and hospitality during our stay in the Kerguelen archipelago.

---

**S1 Fig.**
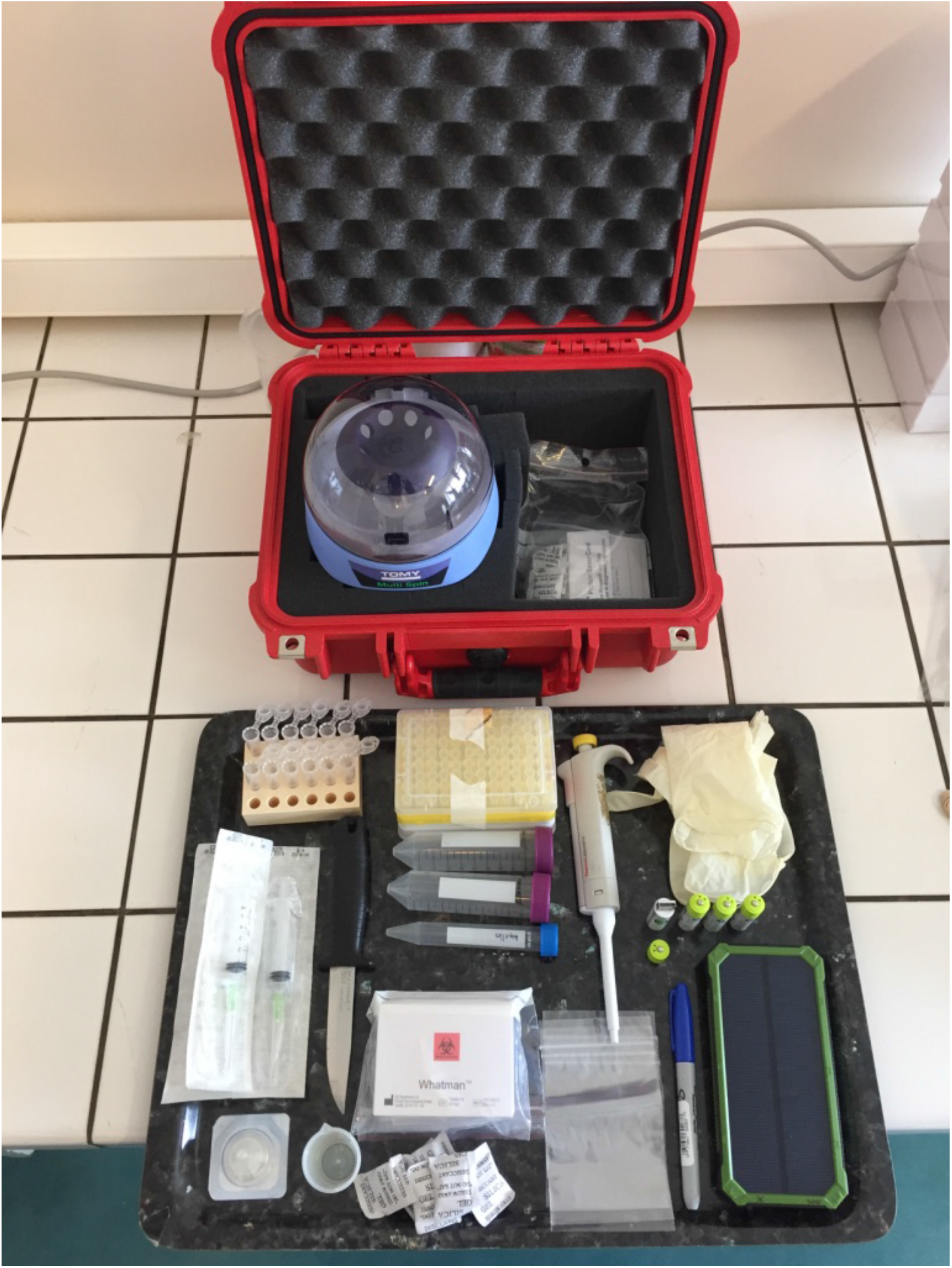
Portable Laboratory carrying case used for liquid biopsy sampling. The case contains a battery powered minicentrifuge and a portable solar panel battery charger for long term expeditions. The case also contains all necessary reagents and equipment to collect biopsies on site.

**S2 Fig.**
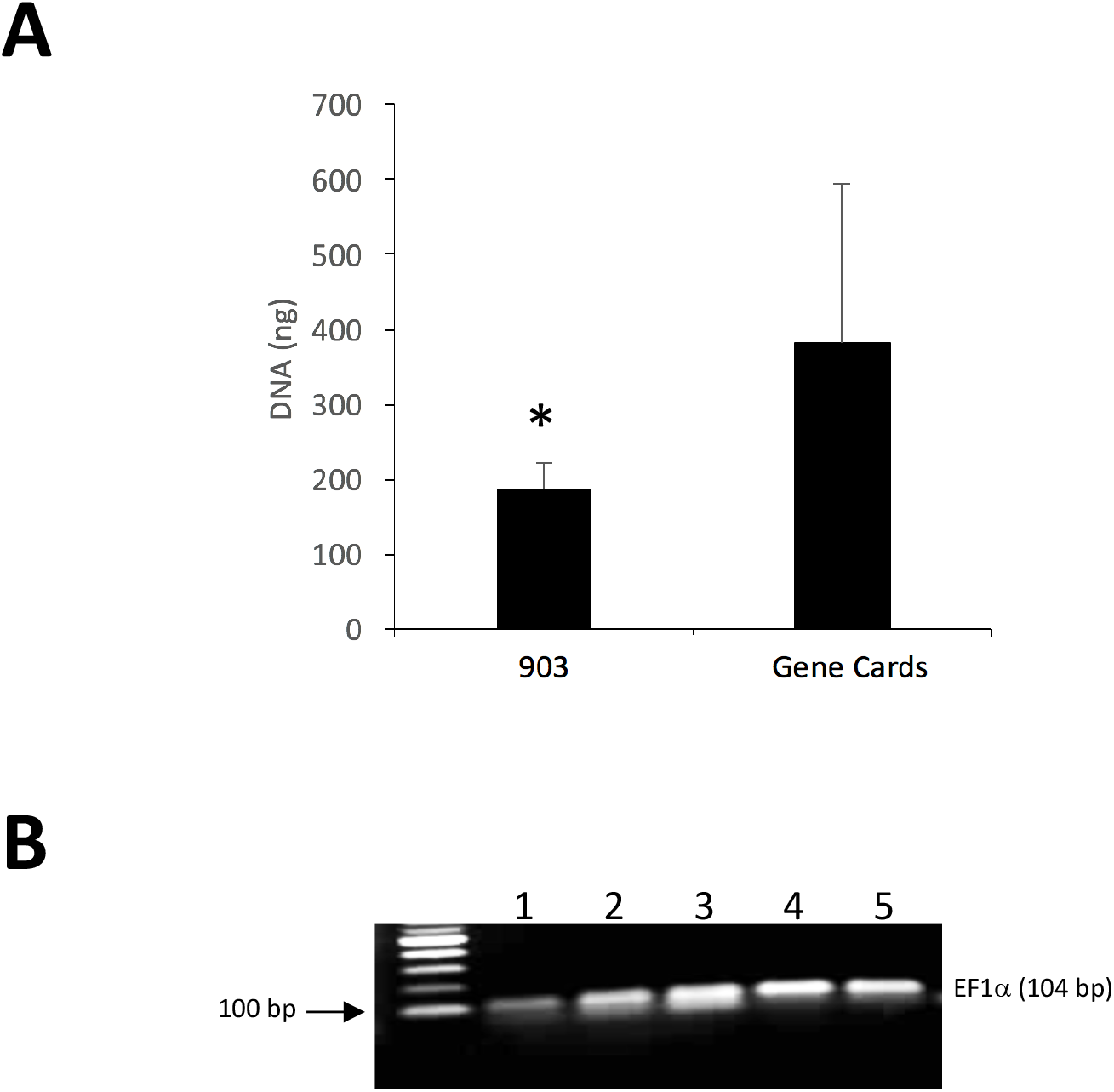
Nucleic acid extraction from FTA cards. (**A**) Comparative analysis of the performance of two different types of FTA cards. DNA recovery from hemocyte pellets that were fixed on the Whatman 903™ cards or the Whatman Gene FTA cards. Values are expressed as the total amount of DNA recovered from a 3 mm diameter sample disc using the PropGEM-based protocol. Results are representative of two independent experiments (n = 12/fgroup). * *p* < 0.05. (**B**) RT-PCR analysis following synthesis of the cDNA template directly from cell pellets fixed on 903 FTA cards without prior mRNA. After the RT step, discs were removed and PCR amplification carried out using ELFα primers. The experiment was carried out using 5 different samples (*M. trossulus*).

## Supplementary Table

**Supplementary Table 1.** RT-qPCR analysis of RNA extracted from frozen or FTA-fixed cell pellets.

| Methodology | Ct | MT |
| --- | --- | --- |
| RNAeasy mini kit | 13,46 ± 0,37 * | 83,38 ± 4,12 * |
| Trizol | 13,14 ± 0,39 * | 84,49 ± 0,05 * |
| Control | 22,89 ± 1,56 | 79,9 ± 0,14 |

Cycle threshold (Ct) and melting temperature (MT) obtained by RT-qPCR for *EF1γ* gene from FTA-fixed RNA extracted using the RNeasy Mini Kit from Qiagen and a standard organic extraction using Trizol extraction protocol. A non-template control (reaction mix with water) was used as control. (\*p-value < 0.05; n= 5 with 3 replicates per sample; data shown as mean ± SD).

